# Bigger Is Better Than Many: A Strategy to Optimize Multi-Gene Co-Expression

**DOI:** 10.1101/2025.09.11.675629

**Authors:** Connor King, Casey-Tyler Berezin, Sarah I. Hernandez, Miranda Nowak, Jean Peccoud

**Affiliations:** School of Biomedical and Chemical Engineering, Colorado State University, Fort Collins, Colorado, 80523, United States of America

**Keywords:** Co-Transfection Efficiency, Plasmid Size, Multi-Gene Expression, Lipofectamine Transfection, Stochastic Modeling, Multi-Plasmid Delivery

## Abstract

Large plasmids are often avoided in mammalian co-transfection due to the assumption that they transfect poorly, driving the use of multiple smaller plasmids. Here, we pair finite-state-projection modeling with flow cytometry experiments to compare one-, two-, and three-plasmid delivery of GFP/BFP/RFP. Estimated entry rates were size-independent from 4.9 to 16.4 kb, indicating that plasmid length is not the dominant barrier in this range. Our results suggest that using lipofectamine slightly increases co-transfection efficiency due to the ability of lipoplexes to contain multiple plasmids. However, this benefit is limited to only delivering two plasmids. Additionally, we show that contrary to current beliefs, putting all genes onto the same plasmid both increases the probability that a cell will express all genes of interest and results in a tighter correlation of gene expression levels compared to these multi-plasmid systems. Together, these results identify multi-cargo delivery and not plasmid size as the key constraint on co-transfection and show that single-plasmid designs are generally preferable for applications such as viral-vector production.

**GRAPHICAL ABSTRACT:** 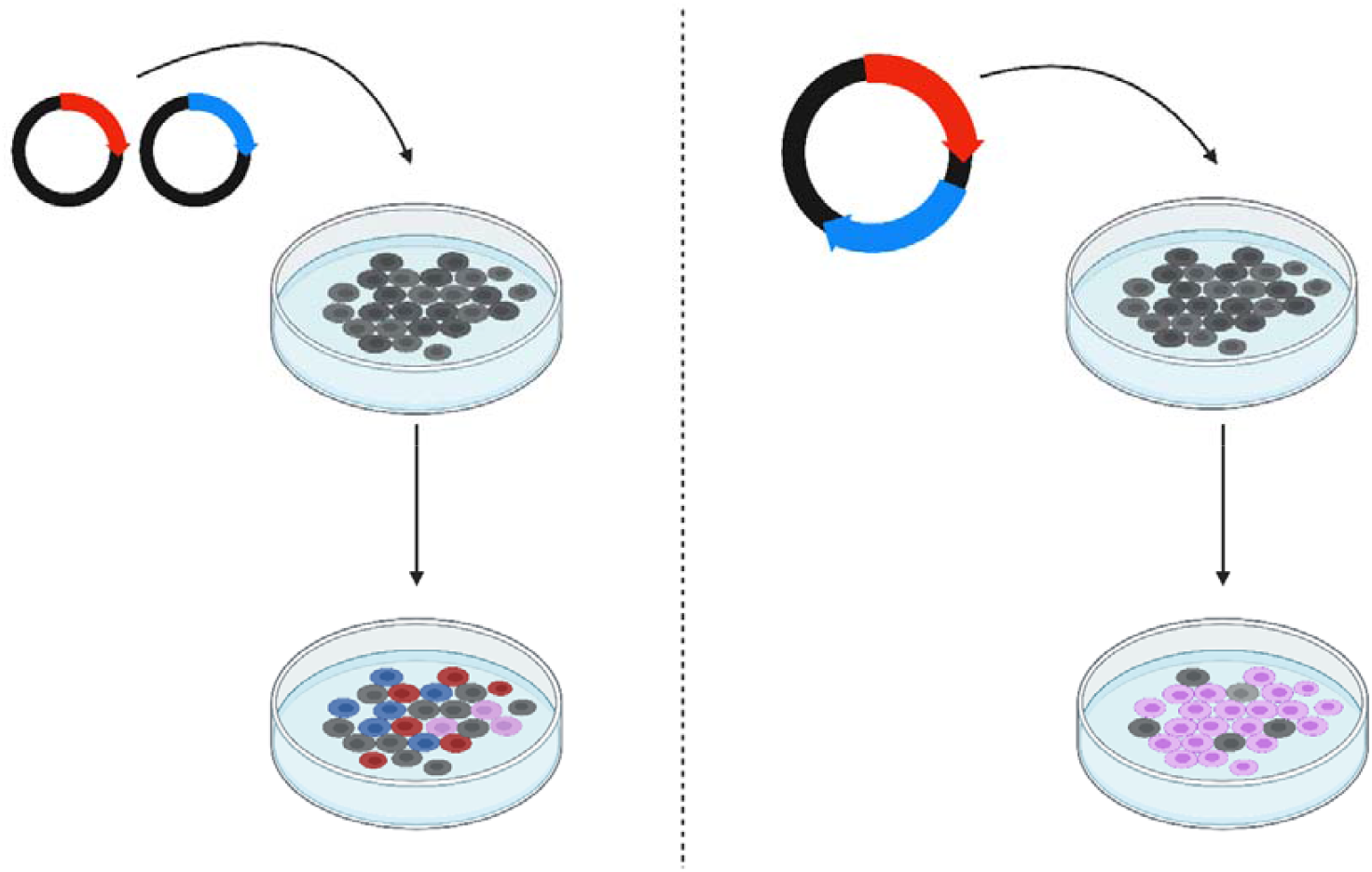

## INTRODUCTION

When producing viruses in mammalian cell lines with plasmids, it is common that multiple helper proteins must be expressed within the cell in addition to the viral genome to produce functional virions. To achieve this, most of these protocols employ the co-transfection of multiple plasmids with each encoding for one these helper genes. This approach stems from a long-held belief that the decrease in transfection efficiency resulting from increasing the size of the plasmid strongly outweighs the decrease in transfection efficiency from delivering multiple smaller plasmids ^1–4^. However, in many cases this leads to very large numbers of plasmids being utilized. For example, Adeno-Associated Virus (AAV), Vesicular Stomatitis Virus (VSV), Ebola Virus, and Influenza B Virus require the simultaneous delivery of 3, 4, 5 and 8 plasmids respectively ^5–9^. These protocols are already challenging as the efficiency of such co-transfections tends to be extremely low. Indeed, this can be seen in the low recovery rates associated with viral production. It begs to question whether this is truly the most efficient way to manufacture these viruses.

There have been several methods proposed for fixing this problem, such as designing cell lines to express some of the genes that might typically be transfected on a plasmid ^5,10^. Another practice that has been proposed is premixing plasmids prior to lipoplex formation as a way to increase the probability of lipoplexes containing multiple plasmids ^11^. However, generating cell lines that stably express these genes is time consuming and may not be possible. Additionally, the increase in co-transfection probability as a result of mixing is rather low (~1.7 fold increase) ^11^. It is critical that we understand the factors that affect co-transfection efficiency and the expression of these genes if we are to design better and more consistent gene expression systems. One method that remains unexplored is whether it is more efficient to put multiple genes onto a few larger plasmids instead of many smaller plasmids, largely due to the general stigma around large plasmids and their perceived inefficiency. Some of this is warranted as there have been many difficulties associated with designing and generating large plasmids historically ^12,13^. However, with improved chemistry becoming more readily available, it is more feasible to produce larger plasmids, warranting the study of their transfection efficiency compared to a multi-plasmid system.

Here we show that it’s possible to make larger plasmids expressing multiple genes without affecting transfection efficiency. We demonstrate that we can consistently achieve higher rates of co-transfection by encoding multiple genes on a single plasmid when compared to 2 and 3 plasmid systems. Additionally, we show that these bigger plasmids have a higher degree of correlation between the level of co-expression of the genes delivered. This ultimately suggests that bigger plasmids could lead to more consistent transfections as well as better control of the level of expression of the respective genes in cells.

## RESULTS

### Plasmid Design

The traditional approach to transfecting mammalian cells with multiple genes involves the co-transfection of multiple plasmids, each with their own expression cassette. However, transfection efficiency decreases significantly as more plasmids are added to the system ^14,15^. Thus, combining multiple expression cassettes into one plasmid can increase transfection efficiency as well as product yield ^16,17^.

We designed bicistronic constructs containing 2 fluorescent protein sequences connected by either a 2A cleavage peptide or an internal ribosome entry site (IRES). Each of these linkers allow coordinated expression of multiple genes; the IRES allows secondary translation initiation of the downstream gene, while genes linked by a 2A peptide are transcribed and translated as one unit and cleaved after translation. The IRES was cloned from a pUC19-T7pro-IRES-EGFP we had previously procured from AddGene (Plasmid #138586). We used a sequence-optimized P2A from VectorBuilder ^18^ and a T2A peptide described in the literature ^19^. Both 2A peptides have a GSG amino acid sequence at the N-terminus to improve cleavage efficiency ^18,20^. For the bicistronic constructs, we introduced mTagBFP2 sequence upstream of the EGFP sequence. For those using a 2A linker peptide, a stopless mTagBFP2 with the stop codon removed was used. We then used these bicistronic constructs to build tricistronic constructs using 2A peptide linkers. We ligated mCyRFP1 downstream of GFP, generating plasmids with BFP-GFP-RFP transcripts.

Several plasmids were designed with shuffled orders of fluorescent proteins to confirm whether the gene order impacted expression levels, notable in the case of the 2A peptide. The two IRES linker plasmids showed decreased expression for the secondary fluorescent protein downstream of the IRES sequences which is consistent with previous findings (data not shown) ^21^. Figure 1 shows some of the different designs for the bicistronic constructs compared to one tricistronic construct, pV58. A full list of the plasmids used in this study are available in Table 1 with Genbank files available in the supplementary data.

**Table 1:** Plasmid designs used in this study.

| Name | Backbone | 5'UTR | CDS 1 | Linker 1 | CDS 2 | Linker 2 | CDS 3 | 3'UTR | Size (bp) |
| --- | --- | --- | --- | --- | --- | --- | --- | --- | --- |
| Unsigned | pUC19 | CMV Pro. | GFP | - | - | - | - | Poly A | 4,966 |
| Signed | pUC19* | CMV Pro. | GFP | - | - | - | - | Poly A | 5,223 |
| pV18 | pUC19* | CMV Pro. | GFP | - | - | - | - | Poly A | 7,287 |
| pV19 | pUC19* | CMV Pro. | GFP | - | - | - | - | Poly A | 12,027 |
| pV20 | pUC19* | CMV Pro. | GFP | - | - | - | - | Poly A | 16,441 |
| pV39 | pUC19 | CMV Pro. | BFP | P2A | GFP | - | - | Poly A | 5,730 |
| pV40 | pUC19 | CMV Pro. | BFP | P2A | GFP | - | - | Poly A | 5,727 |
| pV41 | pUC19 | CMV Pro. | GFP | P2A | BFP | - | - | Poly A | 5,730 |
| pV42 | pUC19 | CMV Pro. | GFP | P2A | BFP | - | - | Poly A | 5,727 |
| pV43 | pUC19 | CMV Pro. | BFP | IRES | GFP | - | - | Poly A | 6,220 |
| pV44 | pUC19 | CMV Pro. | GFP | IRES | BFP | - | - | Poly A | 6,220 |
| pV47 | pUC19 | CMV Pro. | BFP | T2A | GFP | P2A | RFP | Poly A | 6,495 |
| pV48 | pUC19 | CMV Pro. | BFP | T2A | GFP | T2A | RFP | Poly A | 6,492 |
| pV50 | pUC19 | CMV Pro. | BFP | P2A | RFP | T2A | GFP | Poly A | 6,495 |
| pV51 | pUC19 | CMV Pro. | BFP | T2A | RFP | P2A | GFP | Poly A | 6,495 |
| pV58 | pUC19 | CMV Pro. | GFP | P2A | RFP | T2A | BFP | Poly A | 6,495 |
| pV63 | pUC19 | CMV Pro. | RFP | T2A | BFP | P2A | GFP | Poly A | 6,495 |

\*pUC19 backbone with non-functional DNA inserted between origin and ampicillin genes

**Figure 1:**
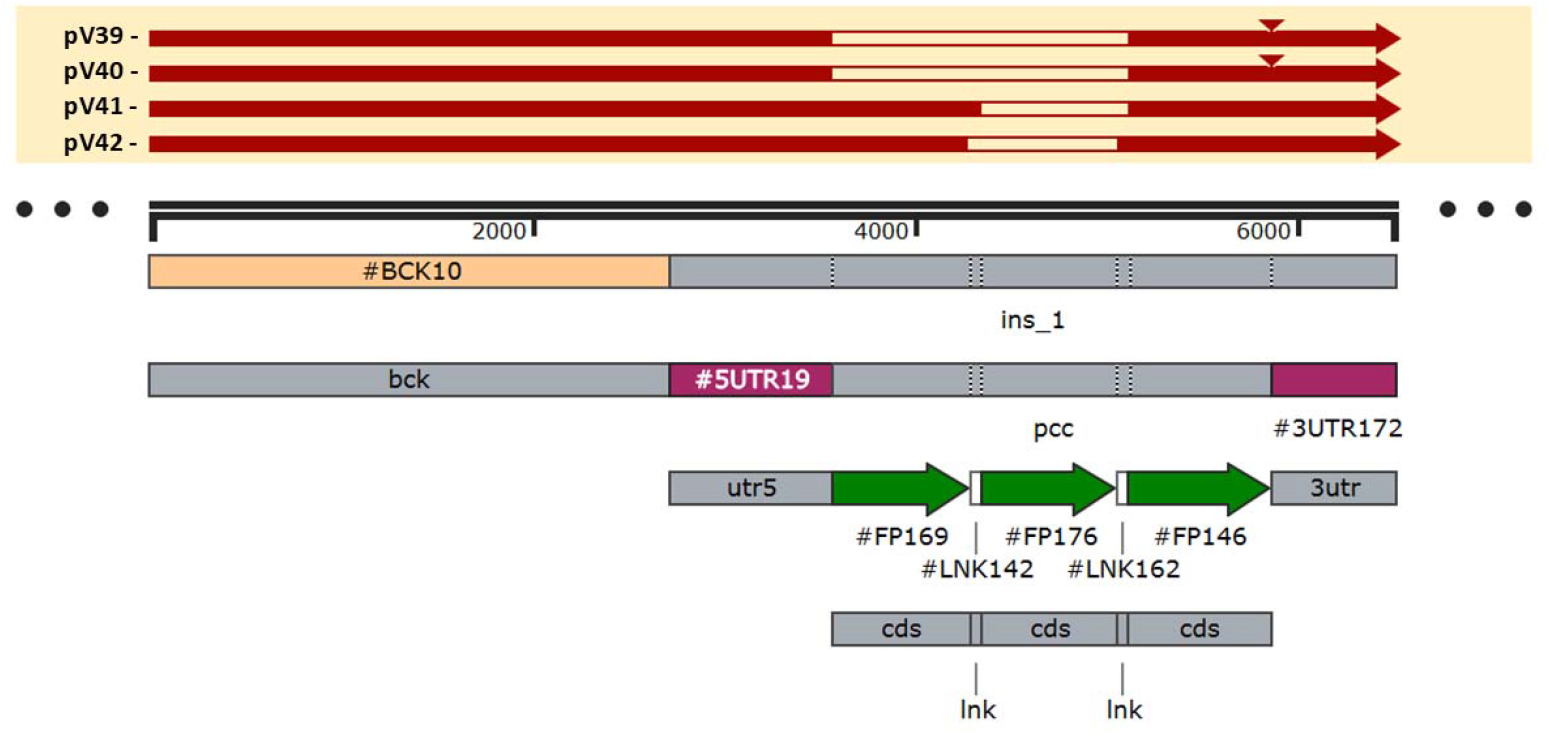
Bicistronic and tricistronic plasmids use shuffled genes to measure expression. The 6495 bp tricistronic plasmid represented by the feature map was constructed with fragments amplified from the bicistronic plasmid with a P2A linker (#LNK142) between a stopless BFP (#FP169), stopless mCYRFP (#FP176) and GFP plasmid with the GFP labeled as (#FP146), using overlap PCR to add in a T2A linker (#LNK162). Several bicistronic plasmids (red) are shown aligned to a tricistronic construct (pV58).

### Model of Plasmid Delivery

We model a single cell exposed to extracellular plasmids (G, R, B). The number of plasmids added is computed from input mass X and length N using the dsDNA relation ^22–24^

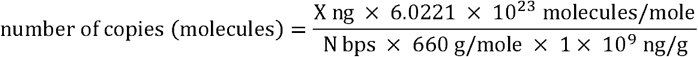

Each plasmid has an entry reaction with rate k × (initial copies). Because lipofectamine complexes can contain multiple plasmids, we compare two models. One where plasmid transfection for each plasmid is independent from the transfection of all other plasmids which we call the “Independent Transfection Model” (Figure 2A). The other model is one where there are additional reactions to account for co-transfection through the same vesicle. This model is called the “Multi-Plasmid Vesicle Transfection Model” (Figure 2B). In literature it has been estimated that each liposome on average contains about 3 plasmids ^11,25^, so we include transfection reactions for each combination of 2 plasmid types and 1 reaction for the entry of all 3 plasmids. Both models effectively only have one parameter and so, log likelihood of observing the data we generated given the model’s predictions was used to evaluate how good the models fit the data.

**Figure 2:**
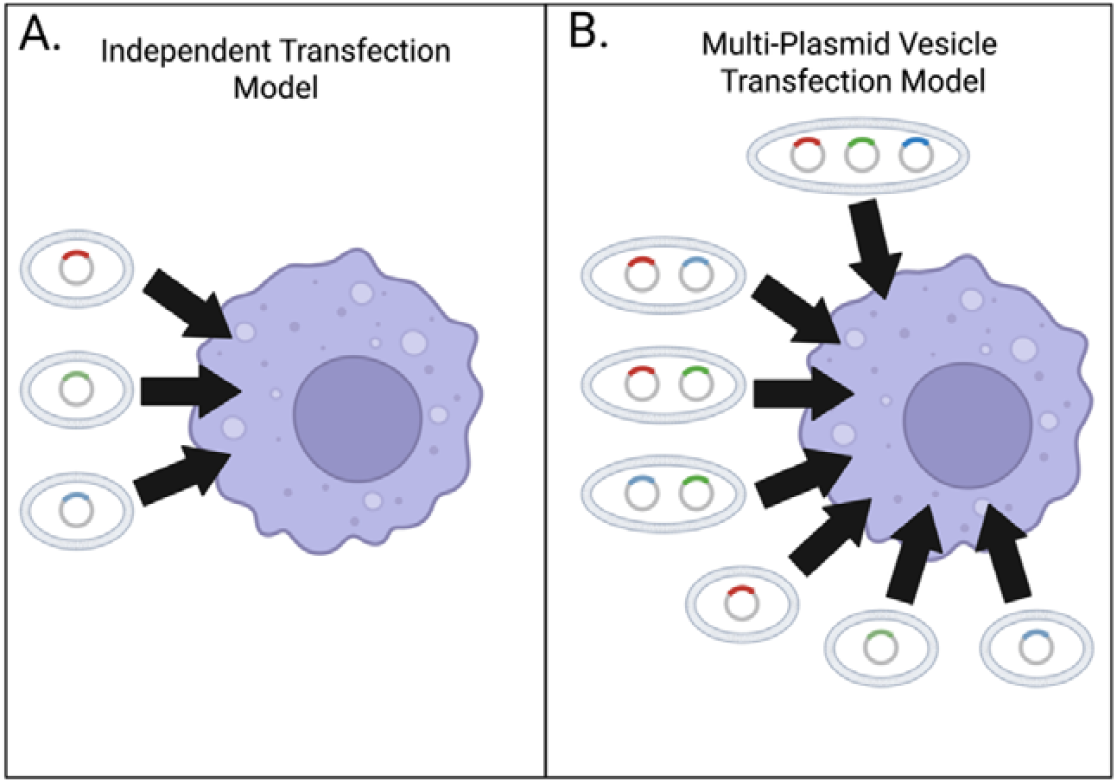
Two Models of Plasmid Transfection via Lipofectamine. (A) The Independent Transfection Model models each vesicle as containing only a single type of plasmid. As such, each plasmid has a single rate of entry that is independent of the other plasmids. (B) The Multi-Plasmid Transfection Model allows for 3 plasmids per vesicle. So, there is an entry reaction for every possible combination of plasmids that can be formed in a vesicle containing 3 plasmids.

To model plasmid co-delivery during transfection, we partition the total entry rate of each plasmid type (G, R, B) into contributions from single, double, and triple co-entry events. These are determined by all possible unordered combinations of 3 plasmids in each vesicle. When all three plasmid types are present, the distribution of vesicle types is: 10% triple-entry, 60% double-entry, shared across the three possible pairwise combinations, 30% single-entry, evenly partitioned among the three plasmids after subtracting co-entry contributions. When only two plasmid types are present, such as G and B, we assume: 2/3 of vesicles are heterotypic and 1/3 are homotypic.

If at least one plasmid of a given type is present in a cell it is considered positive for the fluorescent protein encoded by that plasmid. The rationale for only using the starting number of plasmids in the calculation of entry rates is that the depletion of plasmids from the media due to plasmids entering the cell is negligible. If a single plasmid will cause a cell to be positive and the number of plasmids delivered to individual cells follows a Poisson distribution, the average number of plasmids for all these transfections is less than 1. Considering that the number of plasmids in these transfections are on the scale of 10^6^, the entry of plasmids should not alter the rate of the reaction.

### Plasmid Size Shows no Effect on Transfection Efficiency

The model was initially fit to transfection data from a previous study ^26^ that included GFP-expressing plasmids of varying sizes. The 5 plasmids are 4,966 bp, 5,223 bp, 7,287 bp, 12,027 bp, and 16,441 bp (Table 1). Across all plasmids, the estimated rate constants for cellular entry were highly similar, suggesting no strong size dependence (Figure 3A). This is further demonstrated using linear regression to characterize the relationship between transfection rate constant and plasmid size (Figure 3B). Since the estimated slope of 5.24 × 10^−12^ ± 1.56 × 10^−11^ (p-value =.365, t(3) = 1.066) is not statistically significant, we cannot reject the claim that there is no relationship between plasmid size and plasmid transfection rate. So, for the model predictions in the next sections a single rate constant derived from fitting the model to all the data was used. The value of this universal rate constant was estimated to be 2.22 × 10^−7^ ± 2.48 × 10^−9^ 1/h. The code for fitting the model can be found in Notebook1 on the Github associated with this article.

**Figure 3:**
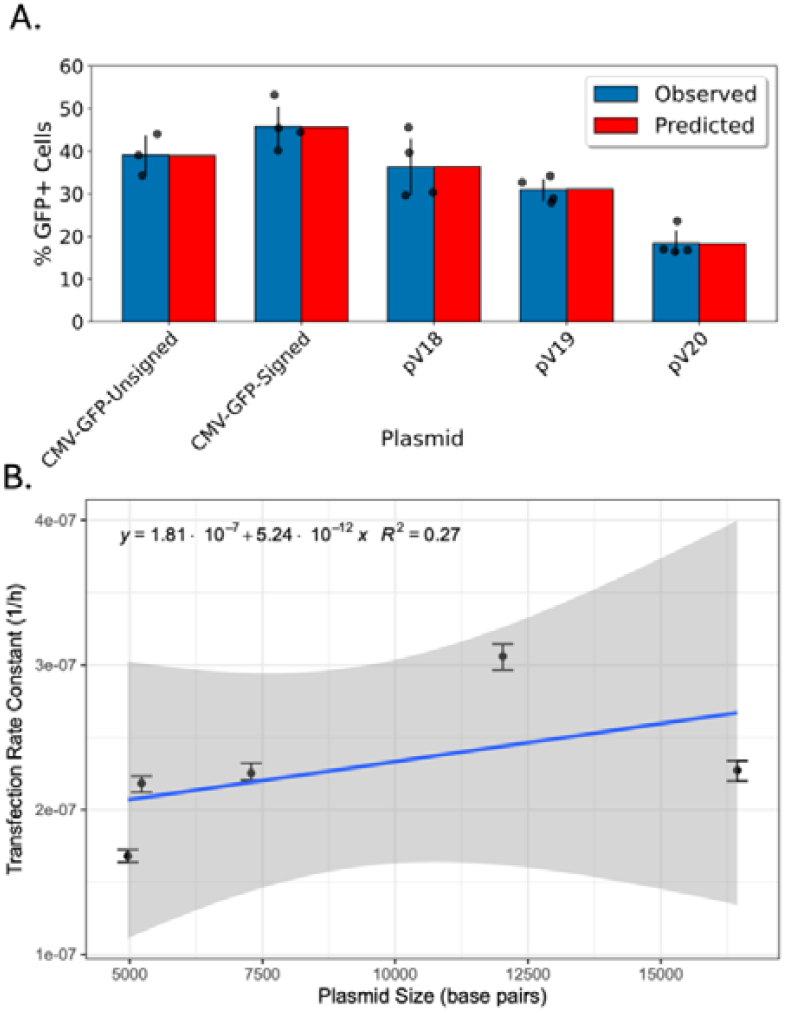
Plasmid Size Shows no Effect on Transfection Efficiency. (A) Observed and fit prediction transfection efficiency of plasmids in lipofectamine transfection experiments from 5 plasmids arranged from smallest to largest. Each dot represents a replicate. Bars show the mean and the corresponding 95% Confidence Interval. (B) Linear regression indicates no significant correlation between plasmid size and entry rate (slope = 5.24 × 10^−12^ ± 1.56 × 10^−11^, t(3) = 1.066, p = 0.365).

### Vesicle Size Explains Co-Transfection Outcomes

Our next goal was to use our models to identify the mode of plasmid delivery by lipofectamine. To achieve this, two triple-transfection conditions were used to examine the effect of plasmid mixing:

- **Pre-mixed:** plasmid solutions were mixed **before** lipoplexes were formed.
- **Post-mixed:** plasmid solutions were mixed **after** lipoplexes were formed, and then the solutions were combined **before** transfection.

Here, we use post-mixing as a control that approximates independent entry of differently labeled vesicles, and pre-mixing to allow co-packaging before vesicle formation. If vesicles routinely carry multiple plasmids, pre-mixing should increase co-expression; if vesicles are cargo-limited and few enter each cell, benefits should be small, especially beyond two plasmids.

Our data suggests that pre-mixing modestly increased double-positive frequency in the two-plasmid system (U = 44.000, p = 0.0496) (Figure S1 A) but had no detectable effect in the three-plasmid system (≥2 positives: U = 88.000, p = 0.0927; all three: U = 15.000, p = 1.0000) (Figure S1 B and C).

This indicates that vesicles co-deliver a limited number of plasmids as this benefit appears to diminish at the triple transfection level. Interestingly, likelihood analyses favored the co-delivery model over independent entry in both pre- and post-mixed samples (pre-mixed −24,422.67 vs −39,769.86; post-mixed −26,085.64 vs −34,885.94) (Figure 4, S2, S3, S4, and S5). This may suggest that despite lipoplexes being formed prior to mixing, there may be some exchange of plasmids between them. The code for simulating these models can be found in Notebook2 on the Github associated with this article.

**Figure 4:**
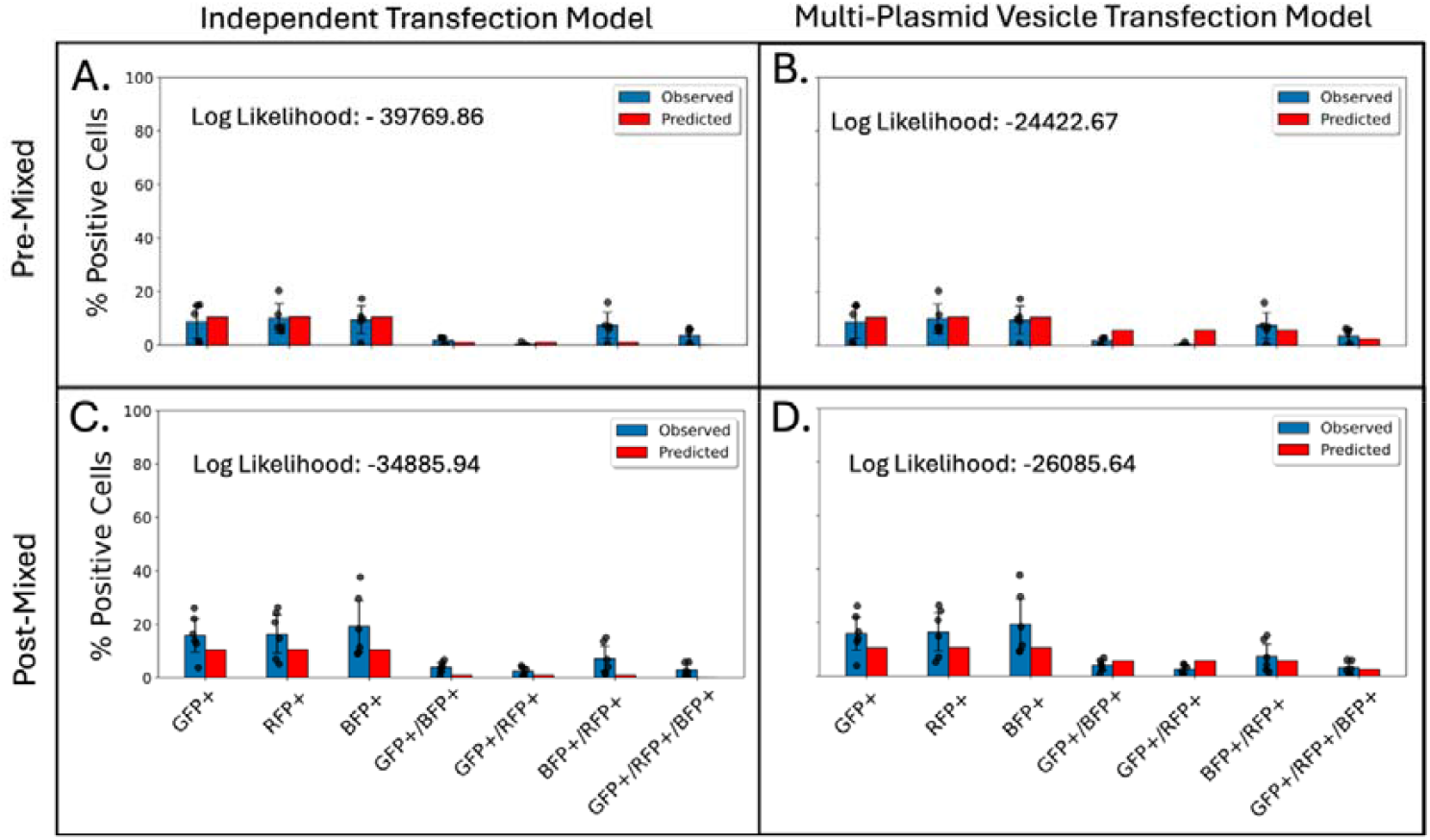
A co-delivery model with a three-plasmid cargo explains observed co-expression patterns. (A). Bar plots show observed versus predicted expression frequencies for triple plasmid transfections under two models. Independent Transfection Model: (A) log likelihoods = –39,769.86 (pre-mixed) and (C) –34,885.94 (post-mixed). Multi-Plasmid Vesicle Transfection Model: log likelihoods =(B) –24,422.67 (pre-mixed) and (D) –26,085.64 (post-mixed). The multi-plasmid model more accurately predicts observed co-expression patterns, particularly in dual- and triple-expression bins. Each dot represents a replicate. Bars show the mean and the corresponding 95% Confidence Interval. Log Likelihood were calculated based on predictions for all 7 combinations of the 3 plasmids.

### Large Plasmids Encoding Multiple Fluorescent Proteins Show Higher Co-Transfection Rates and Co-Expression Correlation than Separate Fluorescent Proteins-Encoding Plasmids

Contrary to common assumptions, our initial findings suggest that combining all fluorescent proteins onto a single plasmid is expected to result in a substantially higher co-transfection efficiency. This expectation arises from two observations: first, increasing plasmid size does not appear to significantly alter the plasmid entry rate constant; second, pre-mixing plasmids in two-plasmid systems leads to only a modest increase in co-transfection efficiency. To test this, we generated six plasmids encoding GFP and BFP, and seven plasmids encoding GFP, BFP, and RFP, and measured their co-transfection efficiency across these three fluorescent proteins.

Our model, using a single transfection parameter, continues to match the experimental data well (Figure S6, Figure S7). While two of the plasmids encoding all three fluorescent proteins deviate from the expected efficiency (Figure S7C, and Figure S7G), the overall trend supports the model’s prediction. The code for simulating the transfection of these plasmids can be found in Notebook3 on the Github associated with this article. Specifically, comparing the probability of dual and triple fluorescent proteins expression between the 1-plasmid systems and the 2- and 3-plasmid systems reveals a statistically significant increase in co-transfection efficiency in the single-plasmid system (Figure 5A, Figure S8; 2-fluorescent proteins expression: U = 281.000, p = 7.88 × 10^−5^; 3-fluorescent proteins expression: U = 114.000, p = 4.88 × 10^−4^).

**Figure 5:**
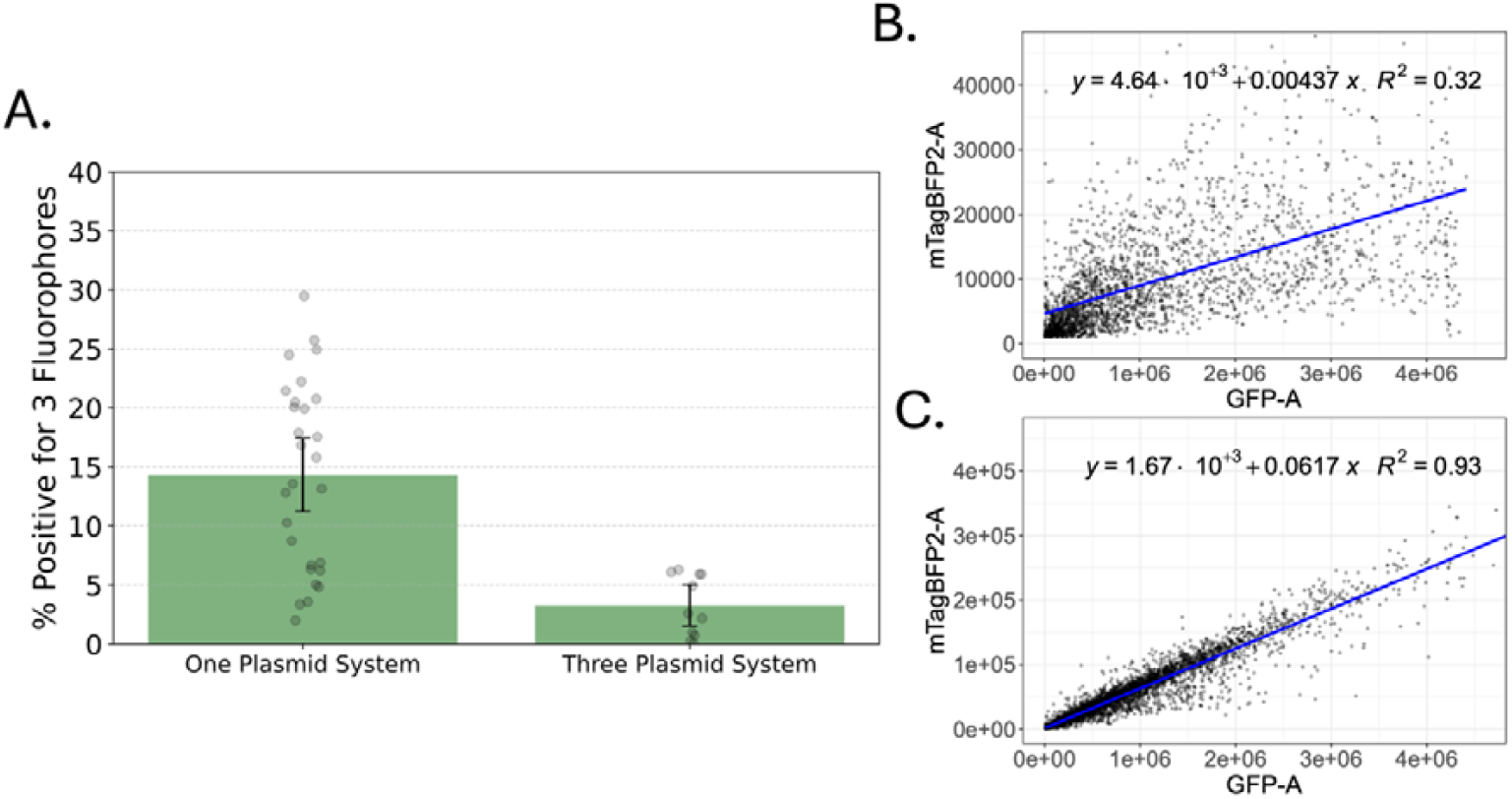
Single-Plasmid Systems Increase Both Co-Transfection Probability and Co-Expression Correlation. (A) Fraction of cells co-expressing two three (GFP+BFP+RFP) fluorescent proteins for single-plasmid (pV47) vs 3-plasmid delivery. Encoding all fluorescent proteins on one plasmid yields a significantly higher probability of triple expression (triple U = 114.000, p = 4.88 × 10^−4^). (B–C) Representative expression–expression plots illustrating stronger correlations when fluorescent proteins are delivered on a single plasmid or on pre-mixed plasmids. single-plasmid systems show the most consistent coordination (higher R^2^).

In addition to increased co-transfection probability, it is not unreasonable to expect that including all genes on the same plasmid would also result in increased correlation in the level of the genes expressed. Indeed, this prediction is confirmed when we observe the correlation between the fluorescent signals observed in the cells. While there are statistically significant correlations observed for both the 3-plasmid system and single-plasmid system, the single-plasmid system consistently exhibits a higher R-squared value (Figure 5B and C, Figure S9 A-C and G-I).

Together, these results indicate that single-plasmid delivery systems may allow for a much consistent transfection and more fine-tuned control over the expression of the genes delivered. It is notable to mention that while there is a correlation observed in expression level in the 3-plasmid system this is only present when the plasmids are pre-mixed. In contrast, the estimated correlations in post-mixed systems were near zero and not statistically significant or negative (Figure S9 D-F). improvement. This along with the weak correlation observed in the pre-mixed system are consistent with our earlier results suggesting that there are multiple plasmids packaged per vesicle, but that the number is modest

## DISCUSSION

This study revisits a foundational assumption in plasmid-based transfection: that increasing plasmid size reduces delivery efficiency. By combining stochastic modeling with flow cytometry experiments across a wide range of plasmid configurations, we demonstrate that this assumption does not hold across the plasmid size range commonly used in mammalian cell systems. Instead, the primary limitation in achieving high co-transfection efficiency arises from the challenge of delivering multiple plasmids to the same cell, not from the size of the plasmid itself. These results are consistent with prior work on the dual promoter expression system ^27^.

We find that transfection efficiency remains effectively constant for plasmids ranging from 4.9 to 16.4 kilobases (Figure 3), supporting the use of a single universal entry rate in our model. Moreover, it appears that the lipoplexes can only contain about 3 plasmids (Figure 4). This limitation results in the very modest increase in co-transfection efficiency that can be observed when plasmids are mixed prior to lipoplex formation in two plasmid systems compared to plasmid mixed post-lipoplex formation, we do not see the same in the three plasmid systems (Figure S1 A) The modest increase in transfection efficiency also is consistent with previous data and models regarding two-plasmid transfection ^11^. Ultimately, the single-plasmid systems consistently outcompete these multi-plasmid transfection systems in terms of co-transfection efficiency (Figure 5A, Figure S8).

Our analysis also reveals that reducing the systems down to a single plasmid also improves the coordination of expression levels among co-transfected genes (Figure 5B and C, Figure S9). Linear regression shows that the single-plasmid systems exhibit significantly stronger expression concordance between fluorescent proteins than either pre or post-mixed systems, which often display weak or inconsistent relationships. This finding highlights a subtle but important advantage of single plasmids as they can reduce variability in relative gene expression.

It is important to acknowledge that these results are for plasmids ranging from 4.9-16.4 kb and may not hold up when considering more extreme plasmid sizes. According to our multi-plasmid transfection model, each vesicle contains about 3 plasmids which is consistent with previous literature estimates ^11,25^. This may suggest that these lipoplexes can contain up to 15 kb of DNA which is rather close to the largest plasmid in this study (16.4 kb). This may be why we do not see any differences in transfection efficiency across plasmids in this study. While they are rare, if we were to utilize plasmid much larger than this, we may see the rate constant begin to decrease due to inefficient lipoplex formation. It should be noted that electroporation, one of the most relevant methods of transfection, does not benefit at all from such packaging effects as there is no formation of complexes in this process (data not shown). So, this methodology would likely benefit strongly from using single-plasmid systems. Additionally, it may be possible that using very small plasmids such as Minicircles could increase the plasmid entry rate constant ^25,28,29^ or could benefit to a greater extent through pre-mixing plasmids prior to lipoplex formation due to more plasmids being present in the lipoplex. In a similar vein, larger plasmids may benefit even less from these effects. Another option could be to use other Polycation nanocarriers such as PEI (which is already used to generate lentiviruses) ^30,31^.

Regardless, these results are widely relevant for most plasmids used in research settings. While generating large plasmids may have been difficult in the past due to limitations in cloning methodologies, many homology-based techniques (such as Gibson Assembly) make the generation of large plasmids from many fragments much more streamline ^32,33^. Our findings suggest that large plasmids should be investigated further as in most practical scenarios they appear to be the most efficient and reliable strategy for achieving robust multi-gene expression in mammalian cells. Alternative approaches using multiple plasmids may still be justified when modularity, differential expression control, or viral compatibility are essential, but these benefits must be weighed against a substantial decrease in co-transfection efficiency and expression coordination. However, for most synthetic biology and cell engineering applications, plasmid consolidation should be considered the default design strategy.

## MATERIALS AND METHODS

### Reagents

The Zyppy-96 Plasmid MagBead Miniprep Kit was purchased from Zymo Research (Irving, CA, USA, #D4102). The long-read library preparation kit and R10.4 flow cell were purchased from Oxford Nanopore (UK, #SQK-RBK114.96 and #FLO-MIN114). Transfection reagents Lipofectamine™ 2000 and Opti-MEM was purchased from Thermo Fisher (Waltham, MA, #11668027 and #11058021). DMEM cell growth media was purchased from Thermo Fisher (Waltham, MA, #21068028) and the fetal bovine serum was purchased from Fisher Scientific (Waltham, MA, #FB12999102). PCR amplification of fragments was carried out using the Q5® Hot Start High-Fidelity 2X Master Mix purchased from New England Biolabs (Ipswich, MA, #M0515). Gibson assembly was performed using the NEBuilder® HiFi DNA Assembly Master Mix purchased from New England Biolabs (Ipswich, MA, #E2621L).

### Biological Resources

The plasmids used were formed from fragments obtained from Twist Biosciences (San Francisco, CA). The HEK293 cells used for transfection and flow cytometry experiments were derivatives of the original tube purchased from ATCC (Manassas, VA, #CRL-1573<SUP>TM</SUP>).

### Plasmid Preparation and Sequencing

#### Plasmid Construction

The plasmids used in this paper were designed and manufactured in house. These plasmids were made using fragments and vectors sourced from Twist Biosciences (San Francisco, CA). The array of plasmids was given unique identifiers (e.g., pV39). DNA fragments were first amplified using the NEB Q5 Hot Start High Fidelity 2X Master Mix according to the manufacturer’s instructions with 25 cycles and annealing temperatures determined using the NEB Tm calculator tool (https://www.tmcalculator.neb.com/). Post amplification, plasmids were quantified using both a Synergy HTX plate reader for A260/280 measurements and a TapeStation 4200 using the genomic screen tapes and reagents to verify fragment size, purity, and quantity. Plasmid assembly was performed using the NEBuilder HiFi DNA Assembly Master Mix according to the NEBuilder design tool available on the manufacturer’s website (https://www.nebuilder.neb.com/#!/), with the exception of an increased incubation time to one hour. In brief, equal molar equivalents of fragments were added to a tube with the master mix. The ligation solution for each plasmid was transformed into chemically competent E. coli cells and grown overnight.

#### Plasmid Purification

The plasmid DNA was extracted from each of the isolates on an epMotion 5075 TC liquid handler (Eppendorf, Hamburg, DE) using the Zyppy-96 Plasmid MagBead Miniprep Kit (Zymo Research, Irving CA, USA), according to manufacturer’s instructions except for changing to pipet mixing during the lysis and neutralization steps as well as an extended final elution time of 10 minutes. Samples were evaluated on a Synergy LX plate reader to determine the quality and quantity of samples after extraction via miniprep.

#### Oxford Nanopore Sequencing

Post purification, 50 ng of each sample was used for sequencing with the MinION Sequencer (Oxford Nanopore, UK). These sequencing libraries were prepared using the Rapid Barcoding Kit (#SQK-RBK114.96) with the Flow Cell (#FLO-MIN114) according to the manufacturer’s instructions. Samples were run on the MinION with a maximum read length kept of 25 kbp. FASTQ files were generated from the super-high accuracy method of the Dorado basecaller within the MinKNOW software and were used for sequence validation, comparison, and evaluation.

### Transfection Experiments

Flow cytometry data were generated from HEK293 cells transfected with several plasmids encoding for GFP, RFP, and BFP. For each experiment, 5 × 10^4^ cells per well in a 96 well plate were plated 24 hours prior to transfection. Cells were transfected at around 70% confluency and the growth media (DMEM supplemented with 10% v/v fetal bovine serum) was changed 4 hours post-transfection. Each well received a total of 100 ng of plasmid DNA, either as a single plasmid or equally divided among 2 or 3 plasmids, (e.g., two plasmids co-transfected at 50 ng each and three plasmids at 33.33 ng each). Two samples (pV58 and pV63) received approximately 60 ng total DNA per well due to limited plasmid stock.

Transfections were performed using Lipofectamine 2000 according to the manufacturer’s protocol using OptiMEM as the incubation media to prepare the lipoplexes. Cells were prepared for flow cytometry analysis 24 hours post-transfection using TrypLE dissociation reagent and PBS. Fluorescent expression was quantified using the Northern Lights spectral flow cytometer to measure 10,000 events per sample. Unmixing was performed with autofluorescence extraction, using unstained cells as a control.

A cell was considered positive for a given fluorescent protein if expression exceeded the detection threshold for that channel. In brief, a positive signal sat outside the defined negative population corresponding to each fluorescent proteins. In the modeling framework, this was defined as the presence of at least one copy of the corresponding plasmid inside the cell. The data unmixing, population gating, and simple statistical analysis were performed using the Cytek SpectroFlo® software version 3.3.0.

### Simulation

The model was simulated using the Finite State Projection (FSP) approach ^34^. The infinitesimal generator matrix was constructed from a defined stoichiometry matrix and a time-dependent propensity function over a finite set of discrete states. Probability mass leaving the truncated state space was tracked via a sink state to ensure normalization. Code implementation followed a previously published structure ^35^. Simulations were run for 4 hours, corresponding to the duration of plasmid exposure in the transfection experiment. The state space was truncated to allow a maximum of eight plasmids per type per cell, based on the assumption that plasmid delivery follows a Poisson distribution. For the fitted rate parameters, fewer than 1 of simulated cells received more than three copies of any plasmid.

### Parameter Estimation

Model parameters were estimated by maximizing the likelihood of the observed flow cytometry data under each model. Posterior uncertainties in parameters were estimated using a Metropolis-Hastings Markov Chain Monte Carlo (MCMC) algorithm ^35–37^. The individual transfection rate constants for plasmid entry rate constant were determined by fitting to transfection efficiency data for GFP-expressing plasmids of varying sizes. The universal plasmid entry rate constant that was ultimately used for the rest of the paper was determined by fitting the model to all the GFP-expressing plasmid data.

### Statistical Analysis

Linear regression was used to assess the relationship between plasmid size and estimated transfection rate constant. Regression models were checked for assumptions, including linearity, homoscedasticity, and normality of residuals. This analysis was conducted using the emmeans and performance packages in R ^38,39^. Mann–Whitney U tests comparing co-transfection efficiencies between pre- and post-mixed plasmid systems as well as 1, 2 and 3 plasmid systems were performed using the mannwhitneyu function from the scipy stats package in Python ^40^. The code for the linear regression analyses performed can be found in the Markdown1 and Markdown2 files on the Github associated with this paper.

## DATA AVAILABILITY

All data generated for this manuscript are publicly available in a repository and can be accessed at https://doi.org/10.6084/m9.figshare.30015583

The code used for this article can be found at https://github.com/Peccoud-Lab/Transfection_Efficiency_Article_2025/

Supplementary Information and Data are available at ACS Synthetic Biology online.

## AUTHOR’S INFORMATION

### Note

J.P. has financial interests in GenoFAB, Inc., a company that may benefit or be perceived as benefiting from this publication.

## AUTHOR CONTRIBUTIONS

CK: Conceptualization, Methodology, Formal Analysis, Visualization, Writing. CTB: Conceptualization, Writing. SH: Data collection, Writing. MN: Data collection.

Jean Peccoud: Conceptualization, Methodology, Writing—review & editing, Funding.

## ACKNOWLEDGMENTS

This work was supported by the National Institutes of Health R01GM147816, and the Suzanne and Walter Scott Foundation. Funding for open access charge: National Institutes of Health.

