## Supplementary Data for "Bigger Is Better Than Many: A Strategy to Optimize Multi-Gene Co-Expression"

**The PDF file includes:**

Figure S1 to S9

Table S1

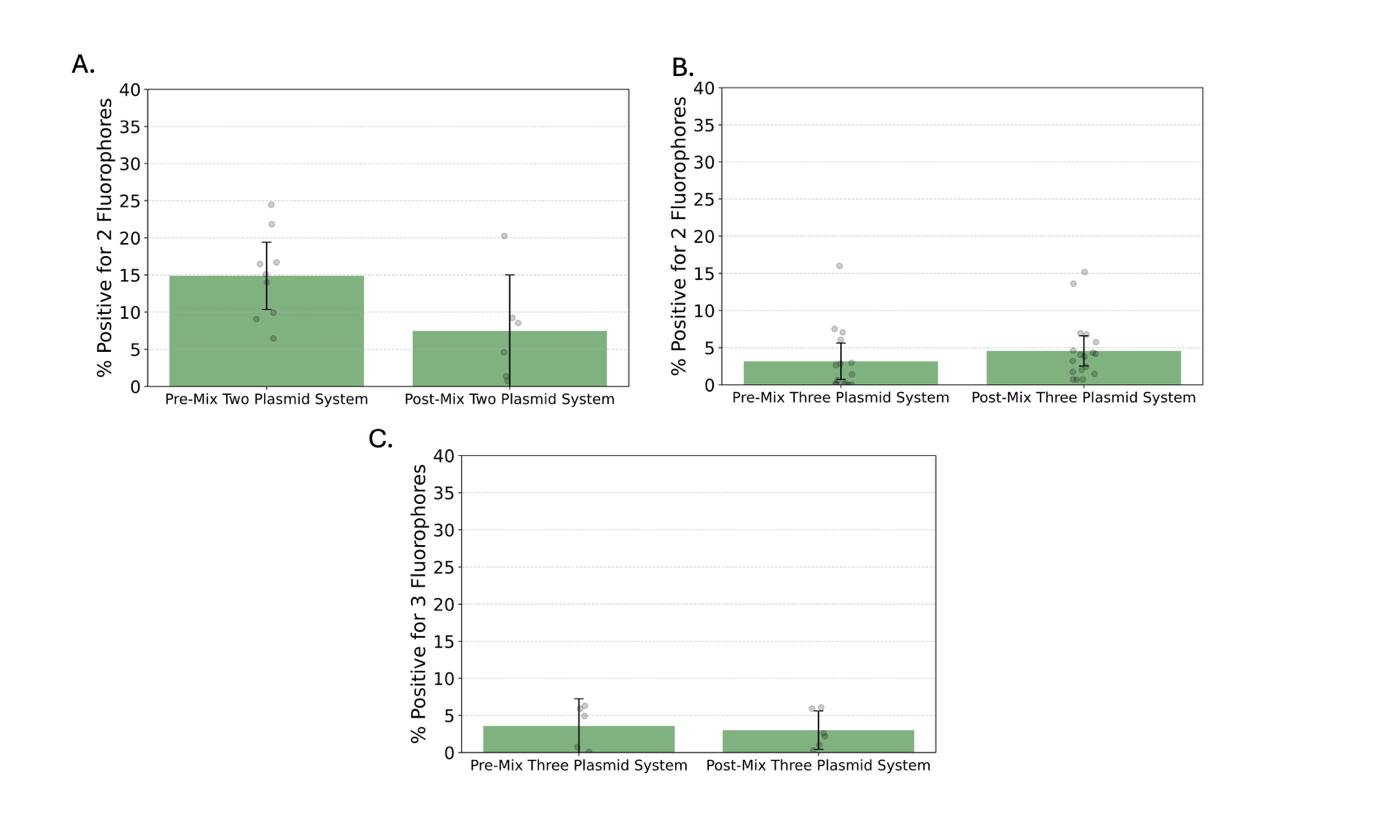

Figure S1. Pre-mixing plasmids improves co-transfection efficiency in two-plasmid systems but not in three-plasmid systems

Bar plots of (A) % of cells positive for two fluorescent proteins in two-plasmid system (B) % of cells positive for two fluorescent proteins in three-plasmid system, and (C) % of cells positive for three fluorescent proteins in three-plasmid system for plasmid samples that were mixed pre or post lipid-plasmid complex formation. The two-plasmid system suggests that pre-mixing increases the percentage of cells expressing two fluorescent proteins (Mann–Whitney U = 44.000, p = 0.0496). However, neither double-positive cells (U = 88.000, p = 0.0927) nor triple-positive cells (U = 15.000, p = 1.0000) show a significant increase due to pre-mixing in the three-plasmid system. Each dot represents a replicate. Bars show the mean and the corresponding 95% Confidence Interval.

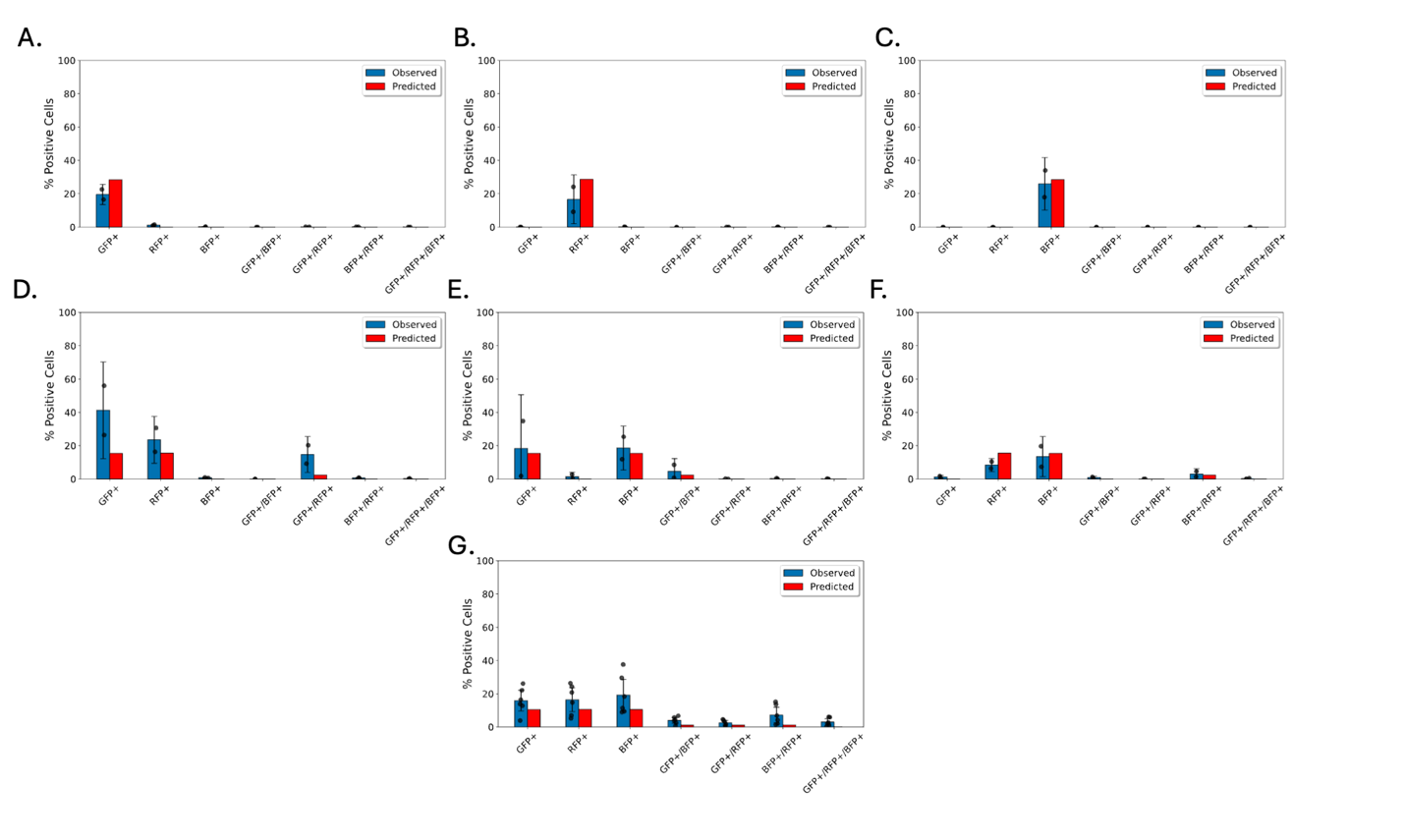

Figure S2. Independent Transfection Model Predictions for Post-Mixed Plasmids

Observed expression frequencies and predicted expression frequencies from the Independent Model for cells transfected with (A-C) one-plasmid, (D-F) two-plasmids, and (G) three plasmids that were mixed post lipoplex formation. Log likelihood = –34,885.94. Entry rate constant used = 2.22 × 10⁻⁷ 1/h.

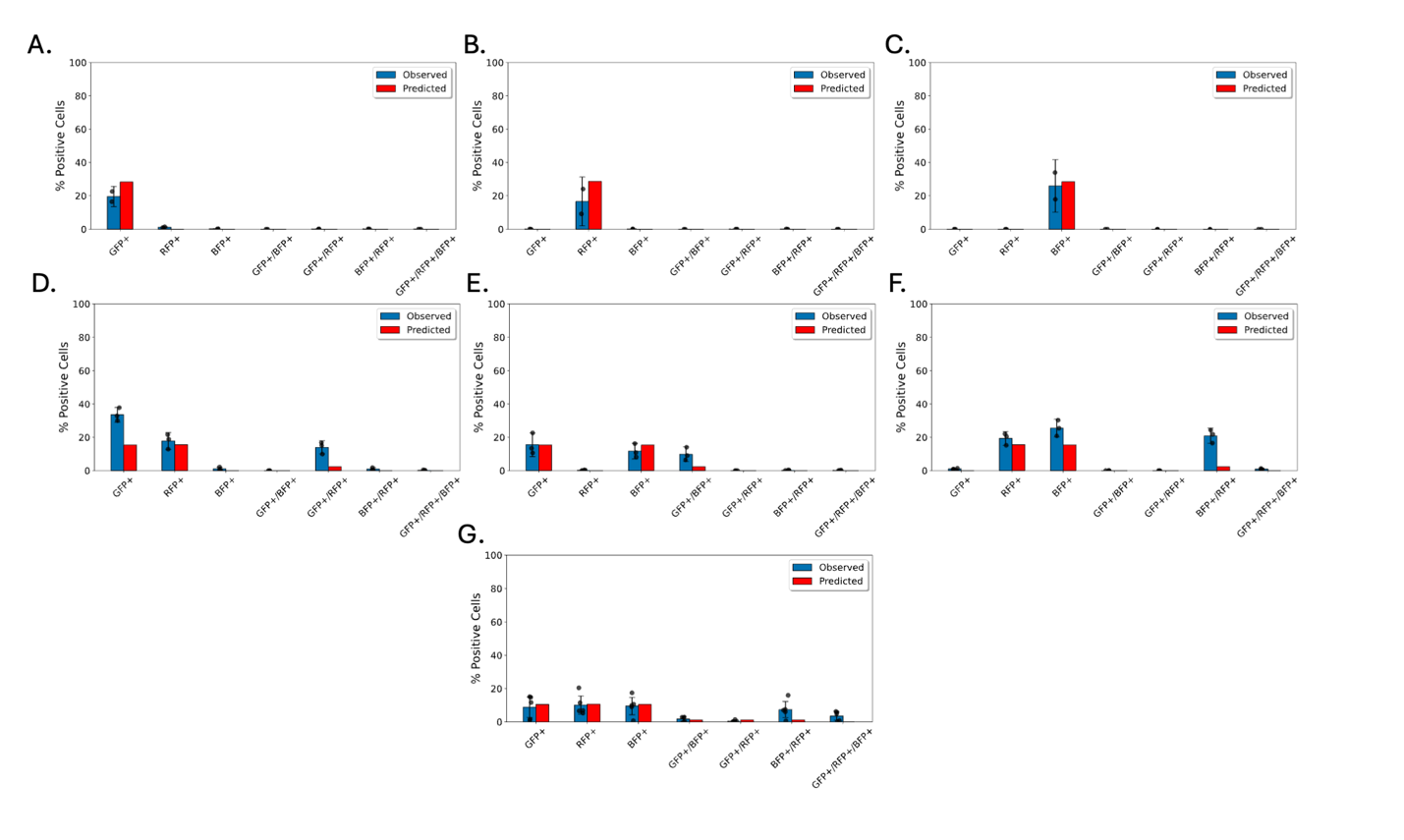

Figure S3. Independent Transfection Model Predictions for Pre-Mixed Plasmids

Observed expression frequencies and predicted expression frequencies from the Independent Model for cells transfected with (A-C) one-plasmid, (D-F) two-plasmids, and (G) three plasmids that were mixed pre lipoplex formation. Log likelihood = –39,769.86. Entry rate constant used = 2.22 × 10⁻⁷ 1/h.

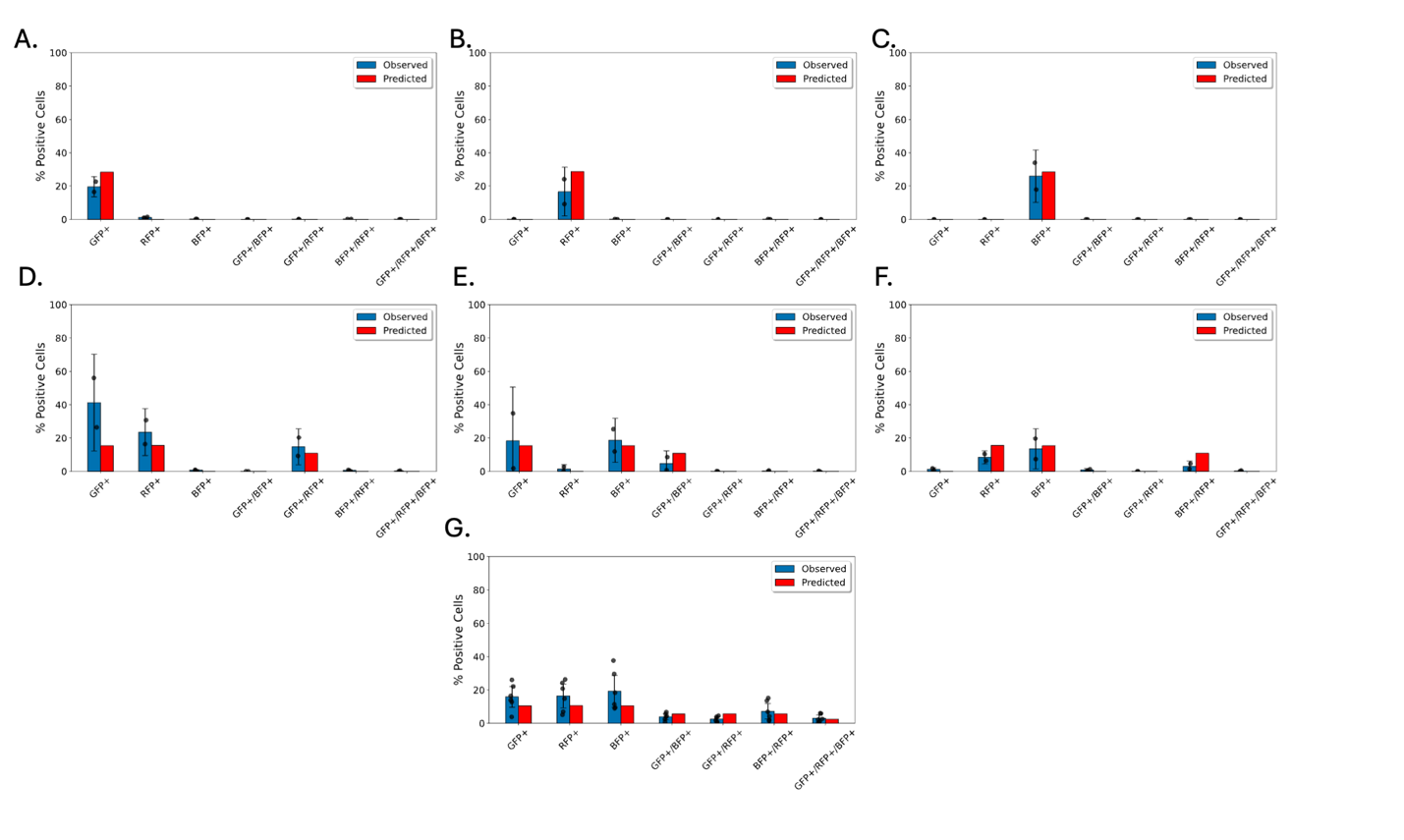

Figure S4. Multi-Plasmid Vesicle Transfection Model Predictions for Post-Mixed Plasmids

Observed expression frequencies and predicted expression frequencies from the Multi-Plasmid Vesicle Transfection for cells transfected with (A-C) one-plasmid, (D-F) two-plasmids, and (G) three plasmids that were mixed post lipoplex formation. Log likelihood = – 26,085.64. Entry rate constant used = 2.22 × 10⁻⁷ 1/h.

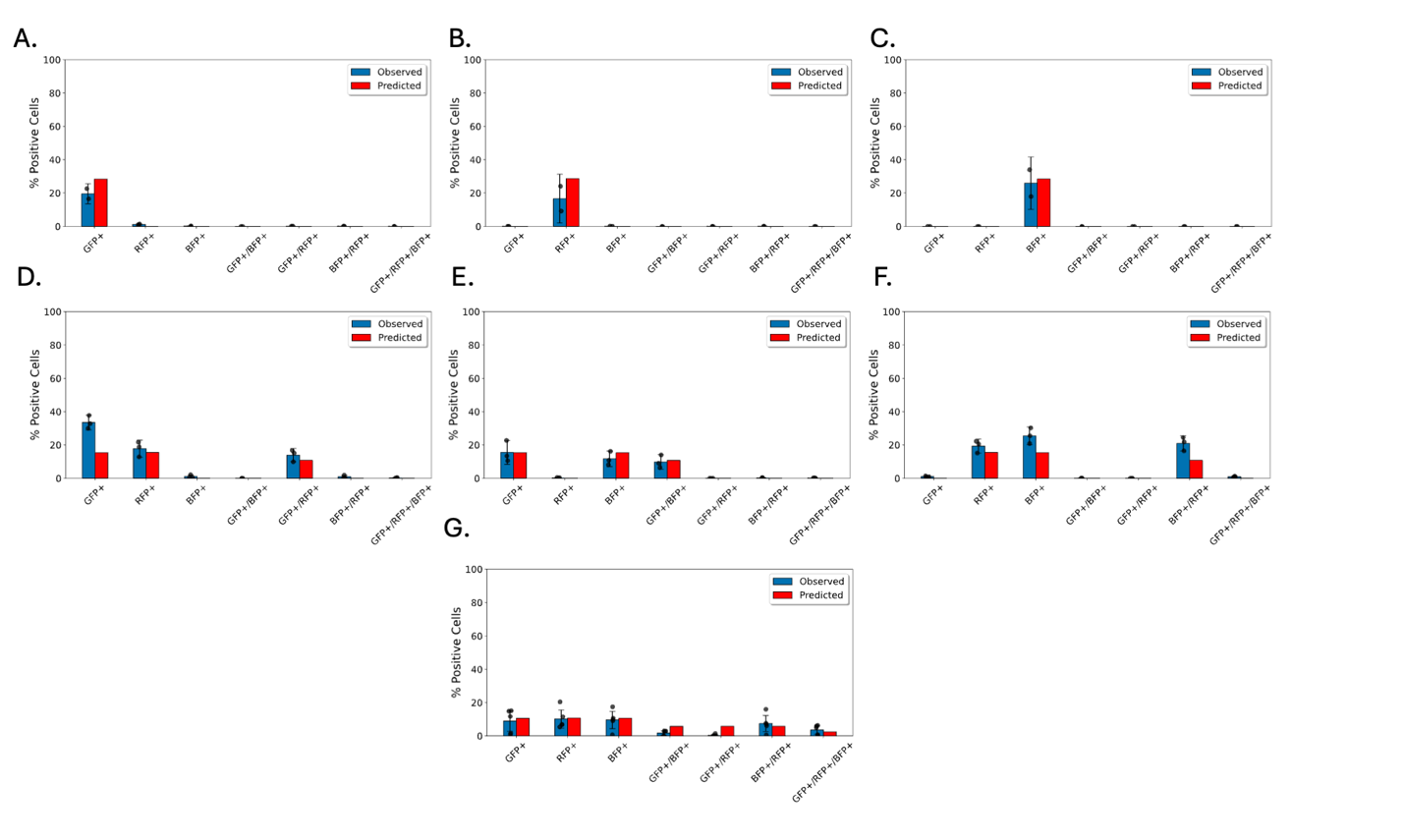

Figure S5. Multi-Plasmid Vesicle Transfection Model Predictions for Pre-Mixed Plasmids

Observed expression frequencies and predicted expression frequencies from the Multi-Plasmid Vesicle Transfection for cells transfected with (A-C) one-plasmid, (D-F) two-plasmids, and (G) three plasmids that were mixed pre lipoplex formation. Log likelihood = – 24,422.67. Entry rate constant used = 2.22 × 10⁻⁷ 1/h.

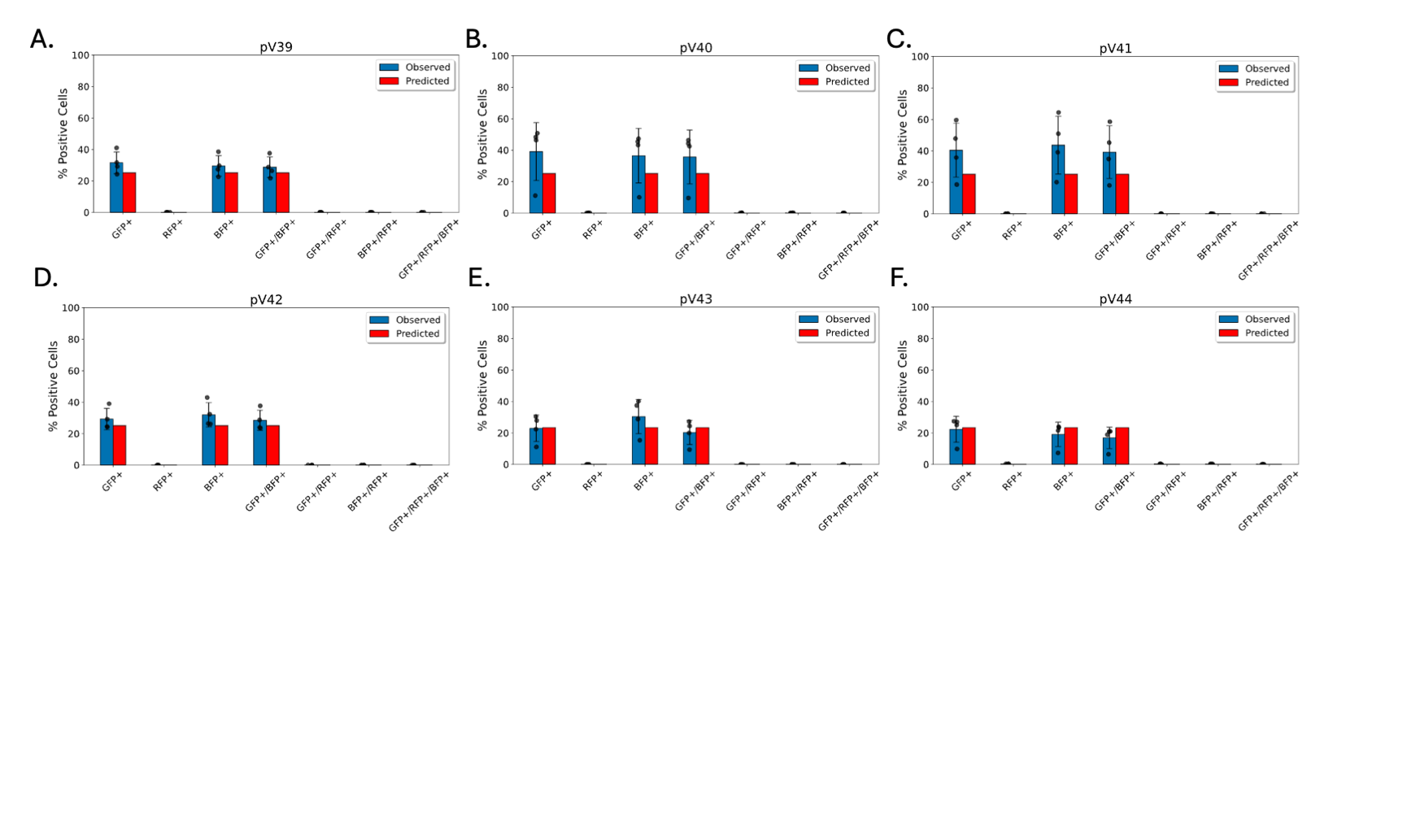

Figure S6. Model Predictions for Two- Fluorescent Proteins Encoding Plasmids

Observed versus predicted frequencies for 6 dual-color plasmids
Observed and predicted expression frequencies for two- fluorescent proteins plasmid constructs. All predictions were made using a single parameter (entry rate constant = 2.22 × 10⁻⁷ 1/h).

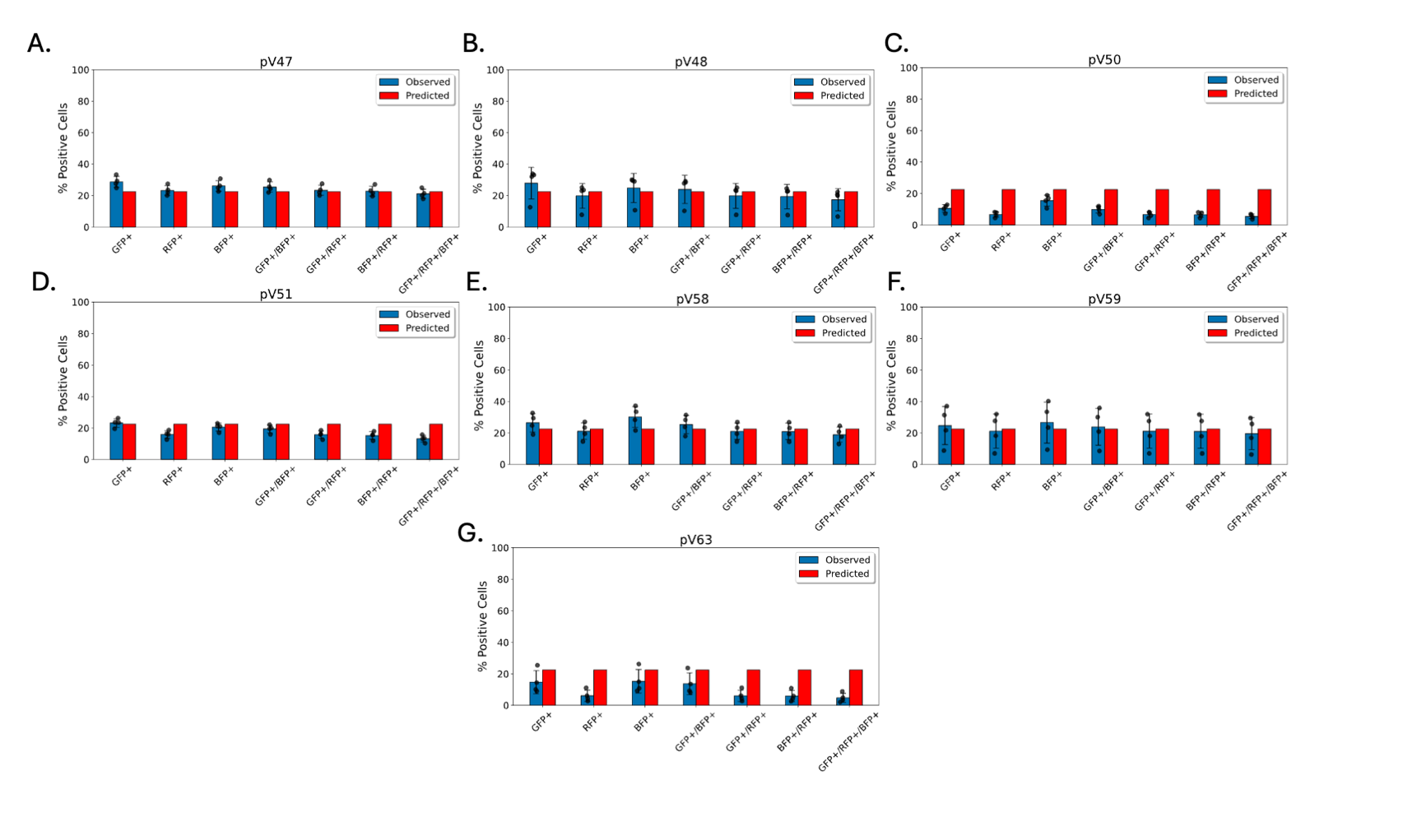

Figure S7. Model Predictions for Three- Fluorescent Proteins Encoding Plasmids

Observed and predicted expression frequencies for three- fluorescent proteins plasmid constructs. All predictions were made using a single parameter (entry rate constant = 2.22 × 10⁻⁷ 1/h).

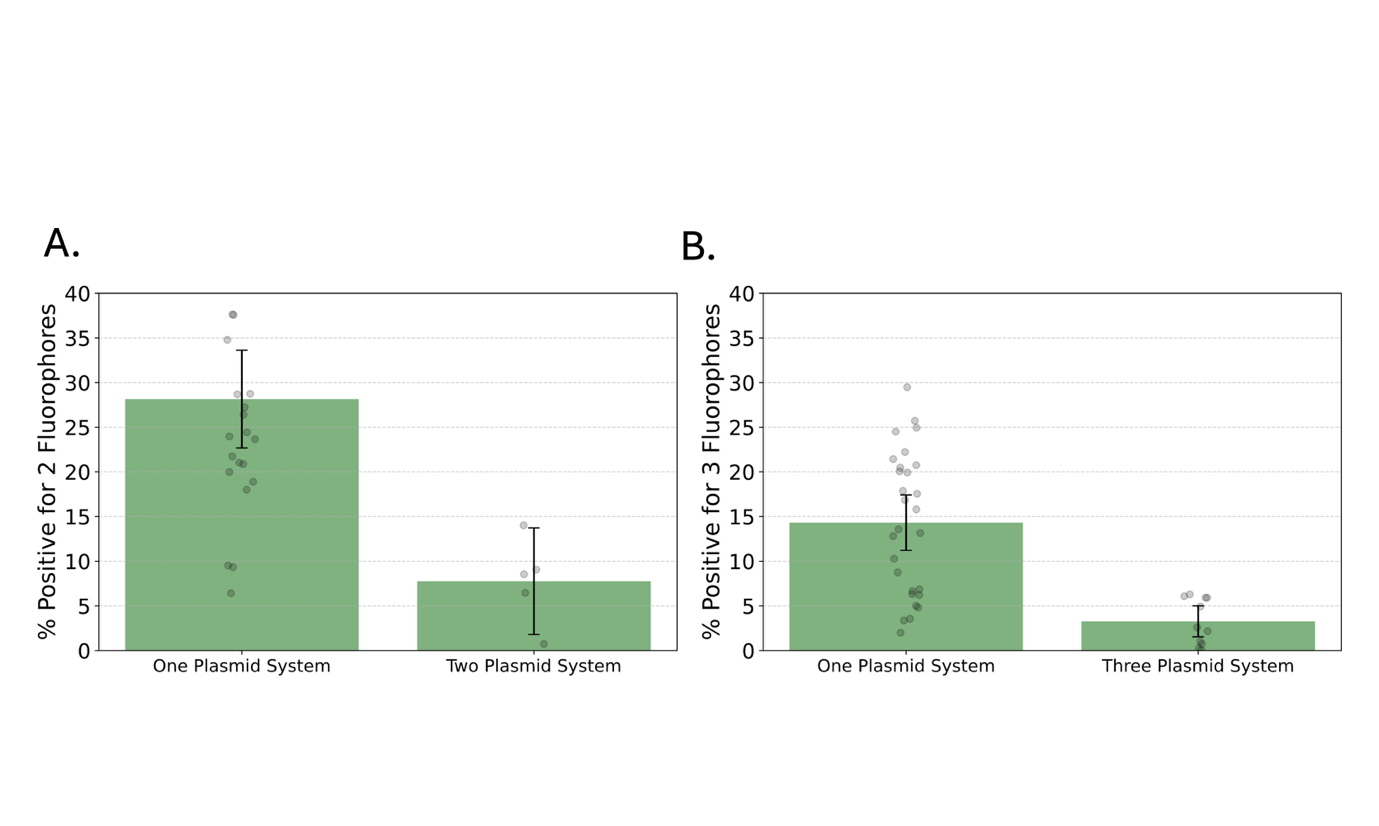

Figure S8 Combining Fluorescent Proteins onto one plasmid increases triple transfection efficiency.

(A) Two- fluorescent proteins co-expression is significantly higher in one-plasmid versus two-plasmid systems (U = 281.000, p = 0.000079). (B) Triple-fluorescent proteins expression is significantly higher in one-plasmid versus three-plasmid systems (U = 114.000, p = 0.0005). Each dot shows an individual replicate; error bars indicate standard deviation. Each dot represents a replicate. Bars show the mean and the corresponding 95% Confidence Interval.

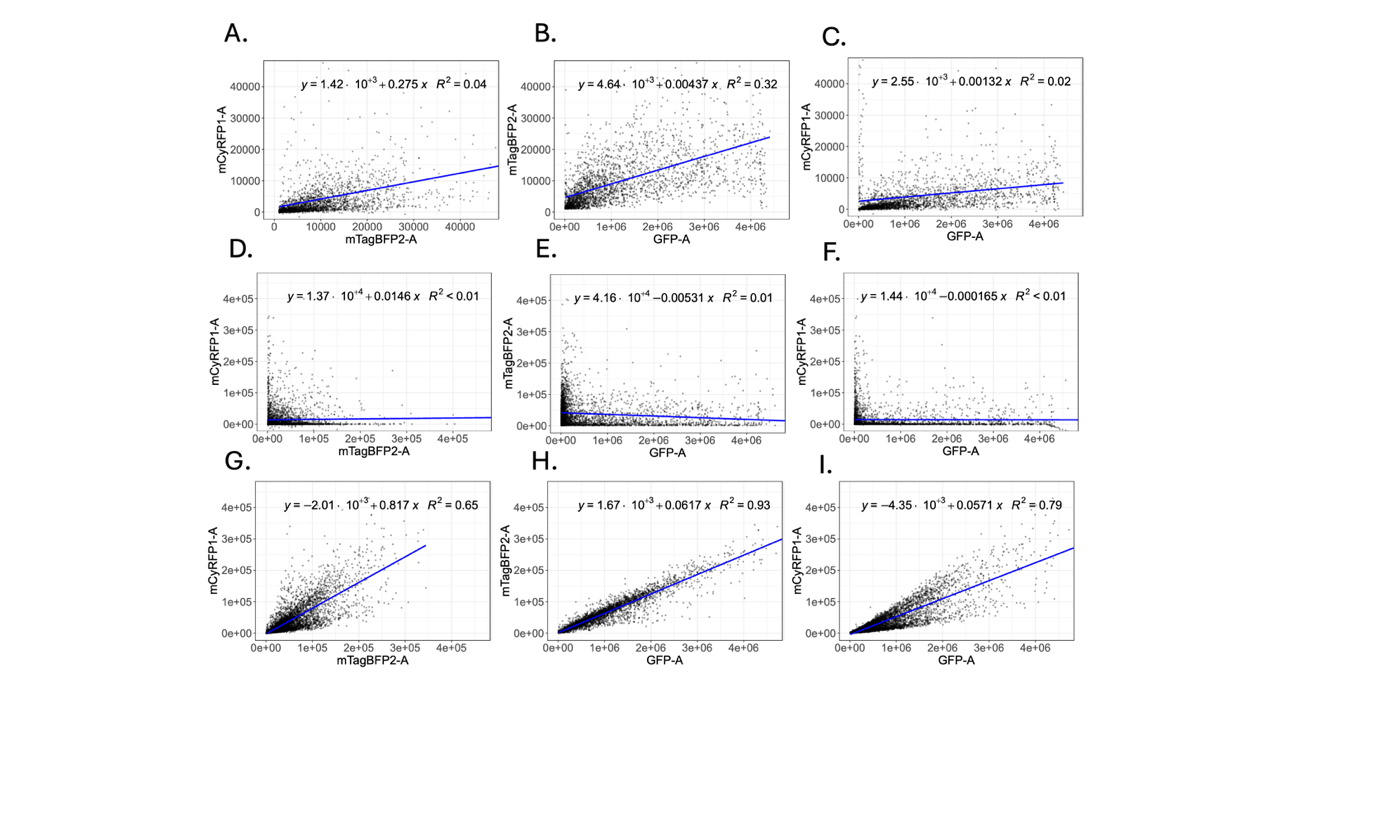

Figure S9

(A) RFP vs GFP expression levels, (B) BFP vs GFP expression levels, (C) RFP vs BFP expression levels in triple positive cells transfected with three plasmids that were mixed before lipoplex formation. (D) RFP vs GFP expression levels, (E) BFP vs GFP expression levels, (F) RFP vs BFP expression levels in triple positive cells transfected with three plasmids that were mixed after lipoplex formation. (G) RFP vs GFP expression levels, (H) BFP vs GFP expression levels, (I) RFP vs BFP in triple positive cells transfected with the single-plasmid system (pV47). The slopes of the best fit lines along with test statistics are found in Table S1.

| Transfection System | Fluorescent Proteins | Slope Estimate | t-test statistic | Degrees of Freedom | p-value | R^2^ |
| --- | --- | --- | --- | --- | --- | --- |
| 3 Plasmids Pre-Mix | RFP vs GFP | 1.32 × 10^-3^ ± 4.13 × 10^-4^ | 6.27 | 2545 | 4.26 × 10^-10^ | 0.04 |
| 3 Plasmids Pre-Mix | BFP vs GFP | 4.37 × 10^-3^ ± 2.48 × 10^-4^ | 34.51 | 2545 | 2.10 × 10^-214^ | 0.32 |
| 3 Plasmids Pre-Mix | RFP vs BFP | 2.75 × 10^-1^ ± 5.27 × 10^-2^ | 10.22 | 2545 | 4.87 × 10^-24^ | 0.02 |
| 3 Plasmids Post-Mix | RFP vs GFP | -1.65 × 10^-4^ ± 1.34 × 10^-3^ | -0.24 | 3368 | 0.81 | <0.01 |
| 3 Plasmids Post-Mix | BFP vs GFP | -5.31 × 10^-3^ ± 1.52 × 10^-3^ | -6.84 | 3368 | 9 × 10^-12^ | 0.01 |
| 3 Plasmids Post-Mix | RFP vs BFP | 1.45 × 10^-2^ ± 2.96 × 10^-2^ | 0.96 | 3368 | 0.33 | <0.01 |
| 1 Plasmid | RFP vs GFP | 6.17 × 10^-2^ ± 3.65 × 10^-2^ | 330.98 | 8125 | << 0.05 | 0.65 |
| 1 Plasmid | BFP vs GFP | 5.71 × 10^-2^ ± 6.50 × 10^-4^ | 172.36 | 8125 | << 0.05 | 0.93 |
| 1 Plasmid | RFP vs BFP | 0.817 ± 1.29 × 10^-2^ | 124.19 | 8125 | << 0.05 | 0.79 |

Table S1: Linear Regression Results for Fluorescent Expression Level Correlation
